# Crohn’s disease patients effectively mobilize peripheral blood stem cells to perform autologous haematopoietic stem cell transplantation

**DOI:** 10.1101/348763

**Authors:** Milton Artur Ruiz, Roberto Luiz Kaiser, Lilian Piron – Ruiz, Tatiana Peña-Arciniegas, Lilian Castiglioni, Priscila Samara Saran, Luiz Gustavo De Quadros, Mikaell Alexandre Gouvea Faria, Rubens Camargo Siqueira, Fernanda Soubhia Liedtke Kaiser, José Francisco Comenalli Marques

## Abstract

**Background:** Treatment with high doses chemotherapy followed by autologous haematopoietic stem cell transplantation is promising for refractory Crohn’s disease patients with no therapeutic option and at imminent risk of further surgeries.

**Objectives:** To evaluate the feasibility and efficacy of haematopoietic progenitor cell mobilization in a group of Crohn’s disease patients preparing for autologous unselected haematopoietic stem cell transplantation in a single institution. This is the first study to evaluate mobilization for Crohn’s disease.

**Methods:** Patients were selected according to criteria of the European Bone Marrow Transplant Society.

**Results:** All patients mobilized with the mean number of haematopoietic progenitor cells obtained and infused being 16.17 × 10^6^/CD34^+^/kg. Most patients required only one leukapheresis session to reach the ideal number of cells. Grafting occurred around ten days after cells infusion. Complications and adverse events during the mobilization period were rare with only one patient presenting sepsis as a relevant event in the period.

Most patients 20 (70%) had anaemia from the beginning of the mobilization but only 11 (37.9%) received packed red blood cell transfusions.

**Conclusion:** Mobilization in patients with Crohn’s disease is effective and it seems they are good mobilizers.

## Introduction

Crohn’s disease (CD), an inflammatory bowel disease of unknown aetiology but with a proven immune component, has an unpredictable, heterogeneous, chronic, and severe course that can affect any part of the digestive tract (1).

It is most commonly observed in the industrialized countries of Western Europe, North America, and both Australia and New Zealand, which were colonized by Europeans. The frequency of the disease is increasing in the 21st century; it is becoming progressively more common in the countries of Eastern Europe, Asia, South America and Africa as a result of the westernization of customs and diet related to globalization (2) (3).

The clinical picture is varied but patients usually evolve with crises of fever, abdominal pain and diarrhoea that are sometimes uncontrollable, in addition to anorexia, emaciation and haematochezia. Anal involvement, which may be the initial manifestation of the disease, is present in about 15-30% of the patients. Fistulas, stenosis and intestinal occlusion make up the complex spectrum of the disease. Extra-intestinal manifestations may occur, as well as the association of CD with other autoimmune diseases (1).

Treatment consists of anti-inflammatory drugs, corticosteroids, immunosuppressants and biological agents, which are used alone or in combinations (4) (5).

Surgical interventions are frequent and it is estimated that more than 50% of patients will have undergone some type of surgical procedure within five years of the onset of the disease. Another point to be emphasized is that after an initial intervention, there is an increased probability of further surgical procedures (1). Thus, when patients are faced with failure to respond to conventional drugs, intolerance or refractoriness to biological agents, and have no other therapeutic option, haematopoietic stem cell transplantation (HSCT) appears as a treatment option.

HSCT is a therapy that has been used to treat autoimmune diseases since the mid-1990s (6). Clinical remissions, both with autologous and allogeneic HSCT in patients with CD concomitant with lymphomas or leukaemias (7) (8) (9), paved the way for the first exclusive procedures aimed at treating the disease (10) (11).

The objective of autologous HSCT is immunological reprogramming with the elimination or reduction of autoreactive lymphocytes and the prospect of inducing a state of tolerogenicity in patients (12) (13).

The procedure begins with a mobilization phase, usually with cyclophosphamide (Cy) and granulocyte-colony stimulating factor (G-CSF), followed by the harvesting of haematopoietic progenitor cells (HPC) from the peripheral blood (PB) that will be re-infused after conditioning with Cy and rabbit or horse anti-thymocyte globulin (ATG). PB-HPC in HSCT with a non-myeloablative conditioning regimen is able to provide hematologic support to reduce the period of neutropenia and thus minimize the risk of infections in this period (14).

## Objectives

To evaluate the results of HPC mobilization in a group of Crohn’s disease patients at a single institution to determine efficacy and safety of the procedure.

## Patients and methods

The criteria consists of active disease (Crohn’s Disease Activity Index above 150, Harvey & Bradshaw score above 4 and Craig Crohn’s Disease Severity score above 17) in addition to the presence of endoscopic evidence of lesions of the digestive tract and refractoriness or intolerance to the use of at least two biological agents. As an additional criterion, the imminent risk of surgery such as rectal amputation or colostomy and non-acceptance of the patient to perform surgery. The mobilization regimen was cyclophosphamide (2 g/m^2^) as a single dose followed by daily filgrastim (10 μg/kg/day) starting five days later until completing haematopoietic progenitor cell harvesting by leukapheresis.

### Ethical considerations

The results described are from patients who received information about the procedure and agreed to the publication of their data by signing a free informed consent form to perform the transplantation procedure. The data are from the project “Autologous Unselected Haematopoietic Stem Cell Transplantation for Crohn’s Disease”, registered in the US platform Clinicaltrials.gov (NCT 03000296). Partial data of this series have been published previously (17).

The authors declare that there are no conflicts of interest.

The criteria for selecting patients for HSCT were those recommended by the EBMT (18) (19) as listed below:

1) Disease activity: Crohn’s Disease Activity Index (CDAI) with a score above 150, Harvey & Bradshaw Index (HBi) above 4, Craig Disease Severity Index (CCSI) above 17;
2) Proven endoscopic lesion of the digestive tract related to the disease and
3) Refractoriness or intolerance to the use of at least two biological agents.

The following was included as an additional criterion:

1. 1) Imminent risk of surgeries including rectal amputation and colostomy implantation and nonacceptance of the patient in respect to surgical treatment.

The data of 29 patients with severe and refractory CD who had been submitted to the procedure between 2013 and December 2017 were analysed in this study. The mononuclear CD34^+^ cell values, day of starting cell harvesting after Cy administration, number of leukapheresis sessions performed, and the day of grafting after cell reinfusion were recorded. Moreover, the clinical parameters of the disease, history and characteristics of the patients prior to HSCT were investigated, as were relevant clinical events and toxicity data observed during mobilization. All results and data were compared to other HSCT studies for CD in respect to the Cy and G-CSF doses, graft composition (selected or unselected), number of cells infused and day of grafting (15) (16).

## Mobilization phase

The mobilization regimen consisted in a single dose of Cy (2 g/m^2^/Kg) followed by G-CSF (filgrastim 10 mcg/kg/day) starting five days after the administration of the Cy and maintained until the end of the harvesting of PB-HPC. The total blood volume processed per session was 20 L in a maximum collection time of six hours.

Leukapheresis sessions were on a daily basis when the number of CD34^+^ cells in the PB was above eight cells/mL. PB-HPC collections was performed using a COBE Spectra RBCX flow path system (MNC program software version 4.7; Terumo BCT, Denver, Colorado, USA).

At the end of the harvesting, patients refrained from major physical activities for seven to ten days before the conditioning phase. More details of the protocol can be find elsewhere (20).

### Statistical analysis

Descriptive statistical analysis was performed from the frequency calculations, central tendency measures and dispersion. In all analyses, a p-value ≤ 0.05 was considered statistically significant.

All data were analysed using the PRISMA statistics program version 6.07 for Windows (GraphPad Software, Lajolla, California, USA, 2015).

## Results

Twenty-nine patients with severe CD refractory to conventional treatments who had been submitted to autologous unselected HSCT were evaluated. Sixteen female and 13 male patients with a mean age of 32.8 years were included. Twenty-one (72.4%) had predominantly stenosing disease in the ileocolonic region, and 22 (75.8%) had a history of disease-related surgical. Eight (27.5%) had perianal involvement and seven (24.13%) had an ostomy at the time of transplantation (table 1).

**Table 1.**
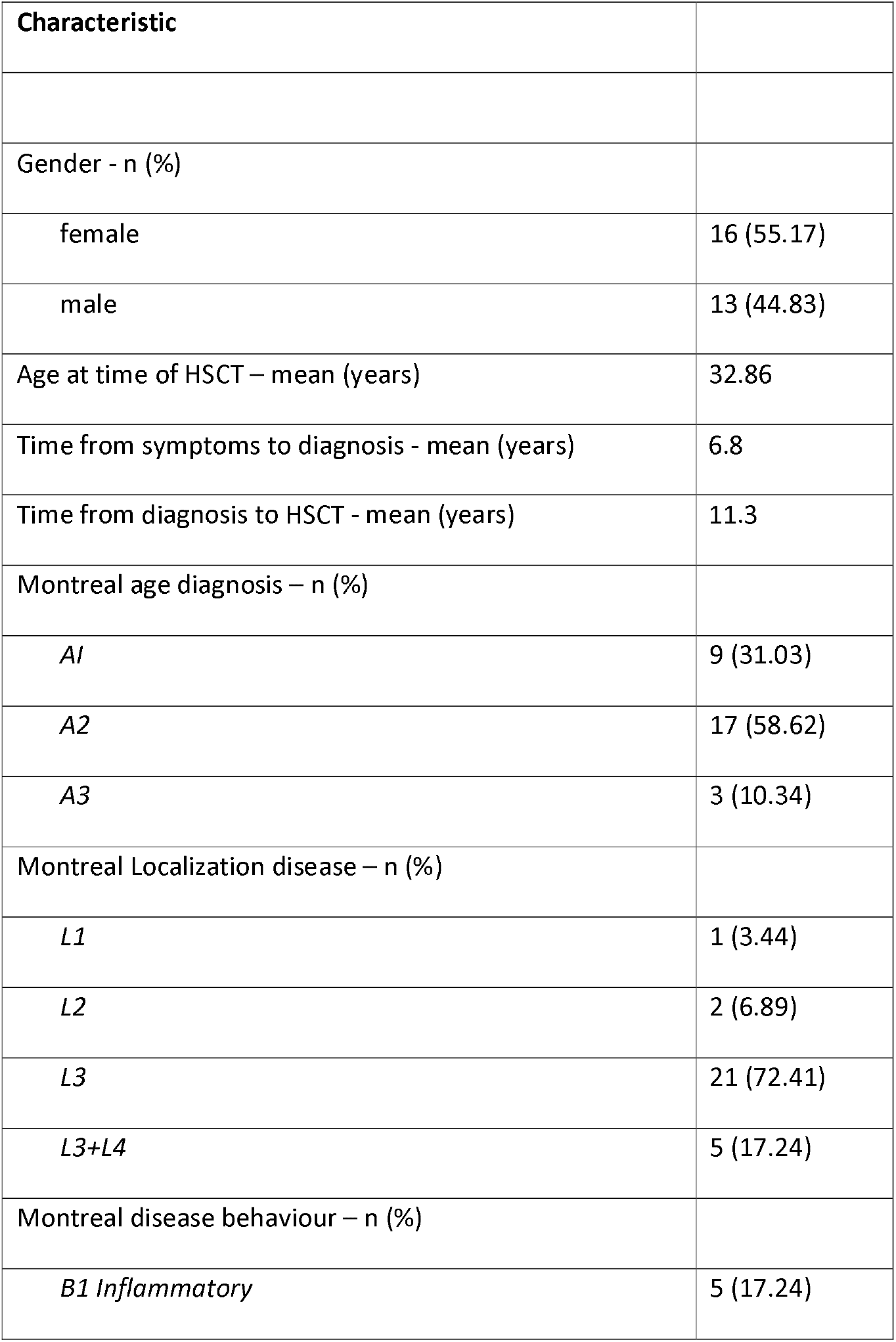
Demographic and baseline data of Crohn’s disease patients prior to haematopoietic stem cell transplantation (n = 29)

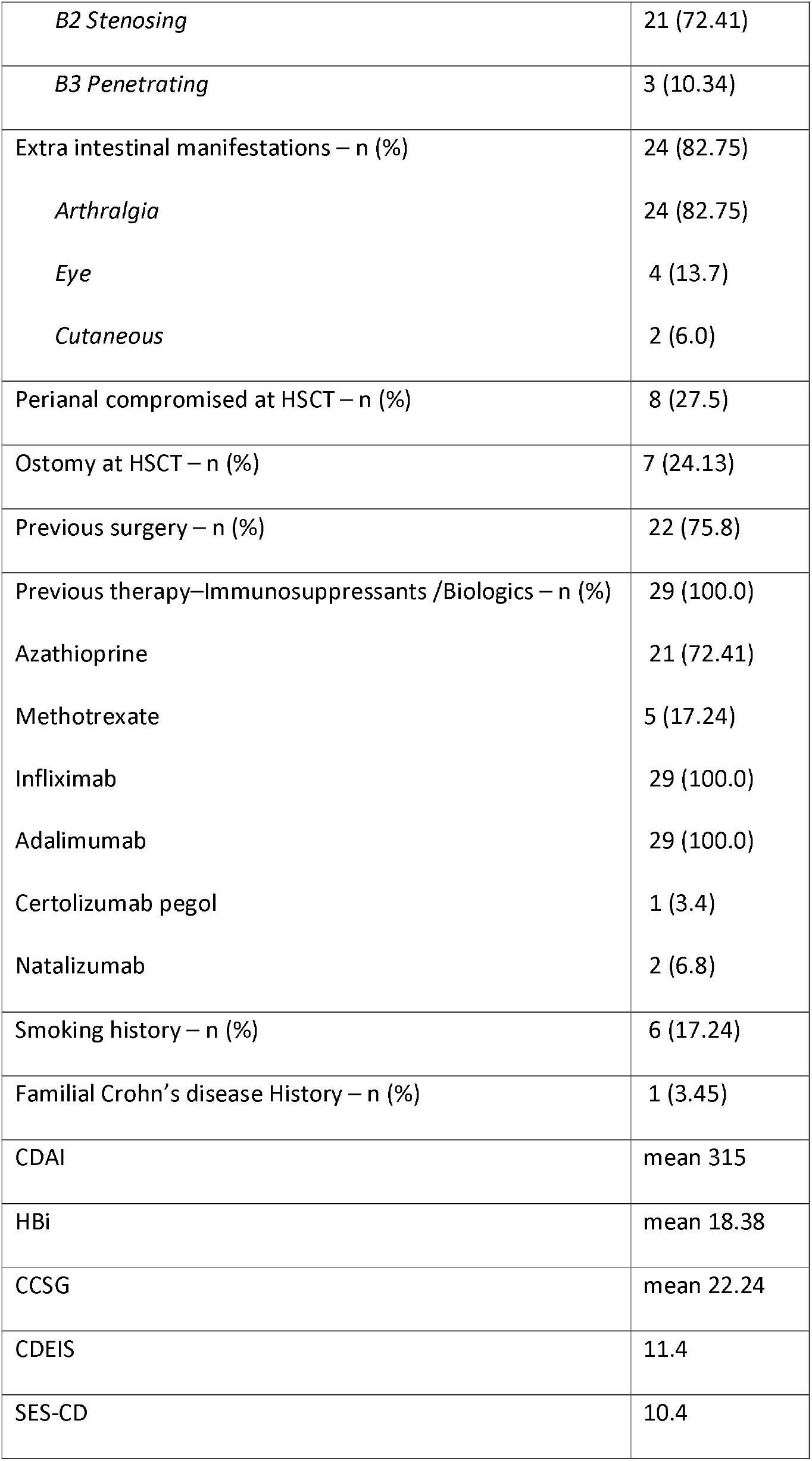

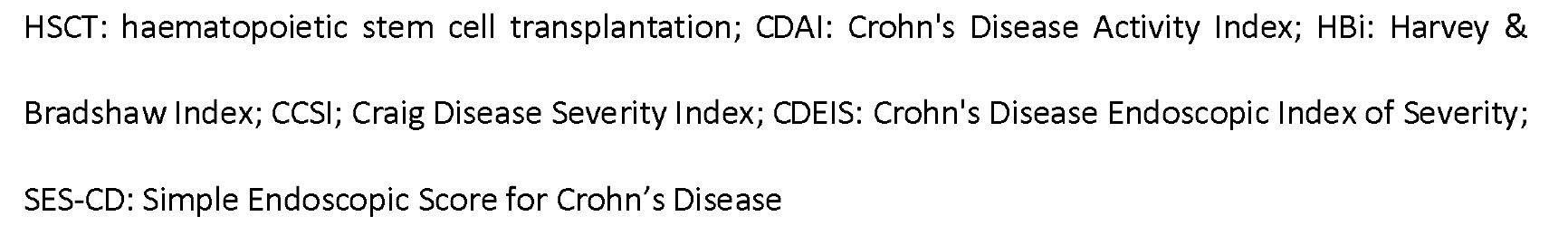

Mean times of 6.8 years and 11.3 years elapsed from the beginning of the symptoms to the diagnosis and from the diagnosis to the HSCT, respectively. The diagnosis was made in the vast majority of the patients (n = 20) after adulthood and the predominant location of the disease was in the ileocolonic region; most patients had stenosing disease.

Most of the patients besides the symptoms in periods of crisis, such as fever, abdominal pain and diarrhoea, presented extraintestinal manifestations with predominance of arthralgia. Other associated autoimmune diseases were not observed in this study, nor were there reports of the necessity of parenteral nutrition despite complaints of anorexia and weight loss.

Of the eight patients with anal involvement, seven were submitted to mobilization for HSCT with ostomies. The majority of the patients had a history of surgeries and a minority were smokers. There were no reports of family cases of CD in this series.

Table 2 shows data on the mobilization and collection of PB-HPC. In most patients (23 - 79.3%), only one leukapheresis session was sufficient to harvest the 3.5 × 10^6^ CD34^+^ cell target. Five of the patients required two sessions while only one patient required three sessions (# 21). The predominant day to start the harvest was Day 10 (62%) with an average of 15.84 CD34 ^+^ cells/kg; patients who started the harvest on Day 11 achieved a mean of 13.81 CD34^+^ cells/kg. The mean overall collection was 16.17 × 10^6^ CD34^+^ cells/kg.

**Table 2.**
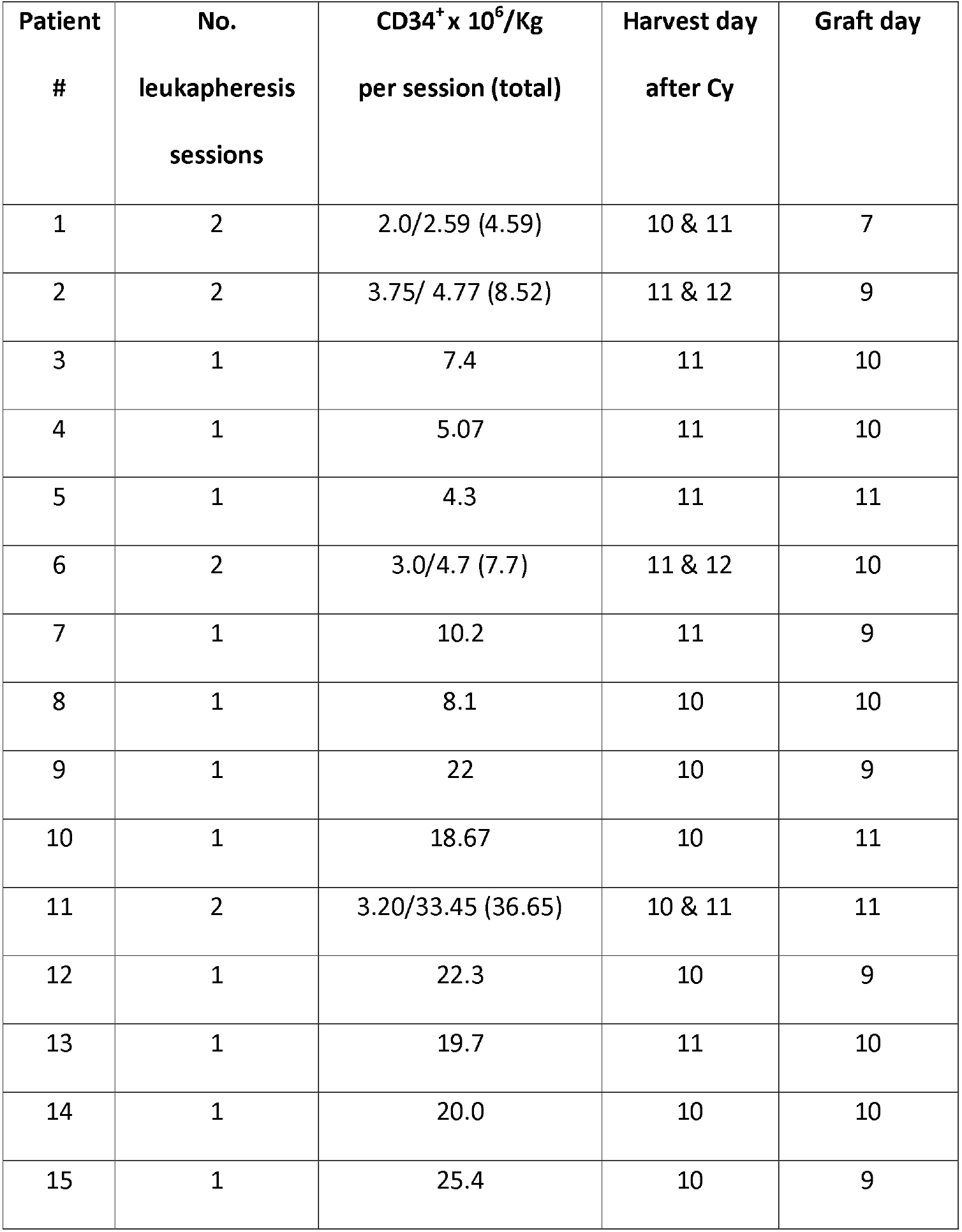
Mobilization Kinetics of haematopoietic stem cell transplantation for Crohn’s disease.

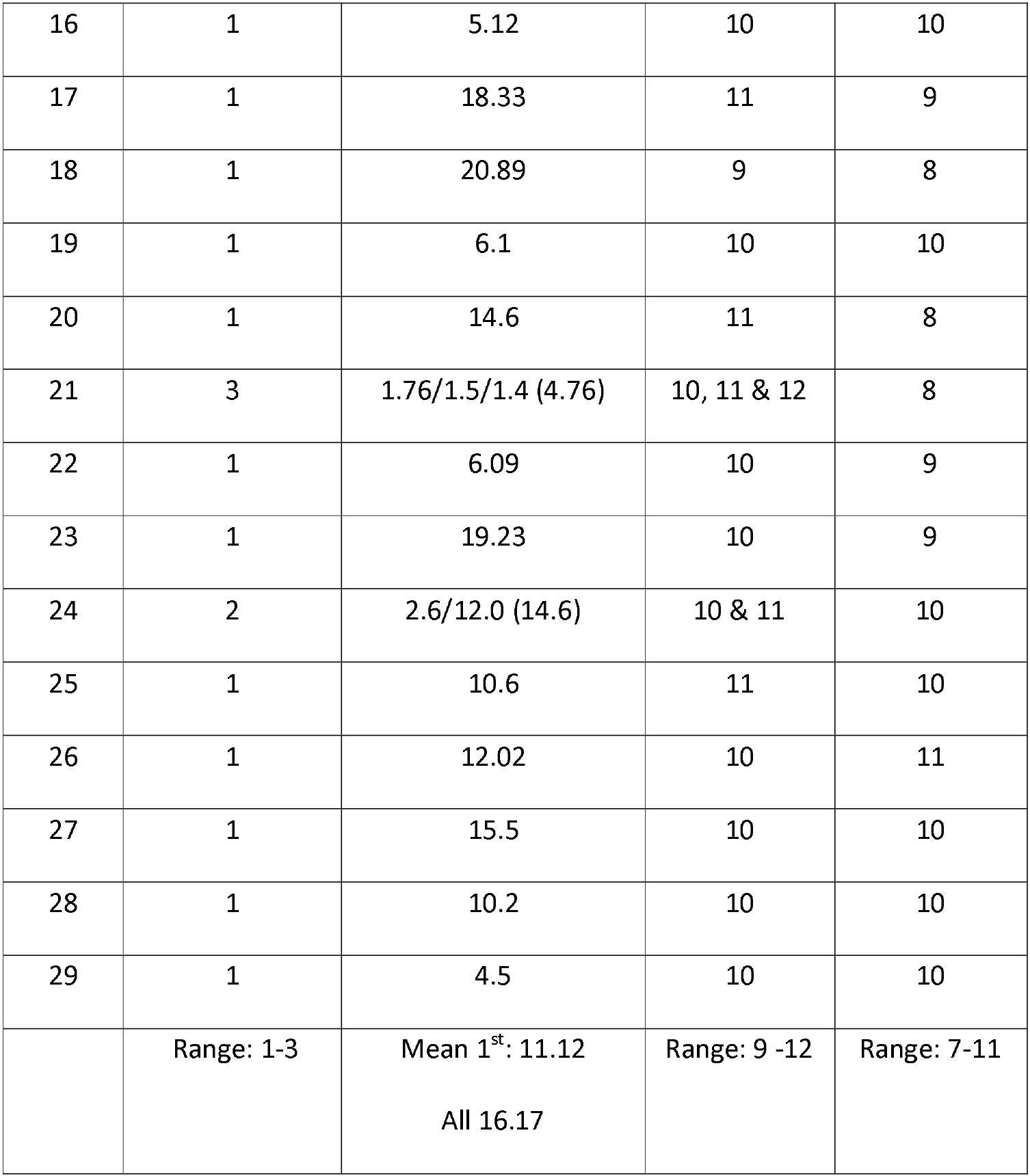

There was no mobilization failure with only three patients presented with marginal collections when the result of the first session was considered in isolation (#1, #21 and #24). The day of grafting occurred on Day 9 (27.5%), Day 10 (45%) or on Day 11 (13.7%) after the reinfusion of the PB-HPC.

Figure 1-a shows a correlation analysis that showed the age of the patient did not influence PB-HPC harvesting (Pearson r: 0.2347; two tailed p-value = 0.2292). Figure 1-b shows that there was no correlation between the time elapsed from the onset of symptoms and the number of PB-HPC harvested (Pearson r: 0.2451; two tailed p-value: 0.2087). Figures 1-c and 1-d shows, using Pearson correlation analysis, that there is no difference in the results of transplanting selected or unselected cells and in the Cy dose used for mobilization.

**Figure 1.**
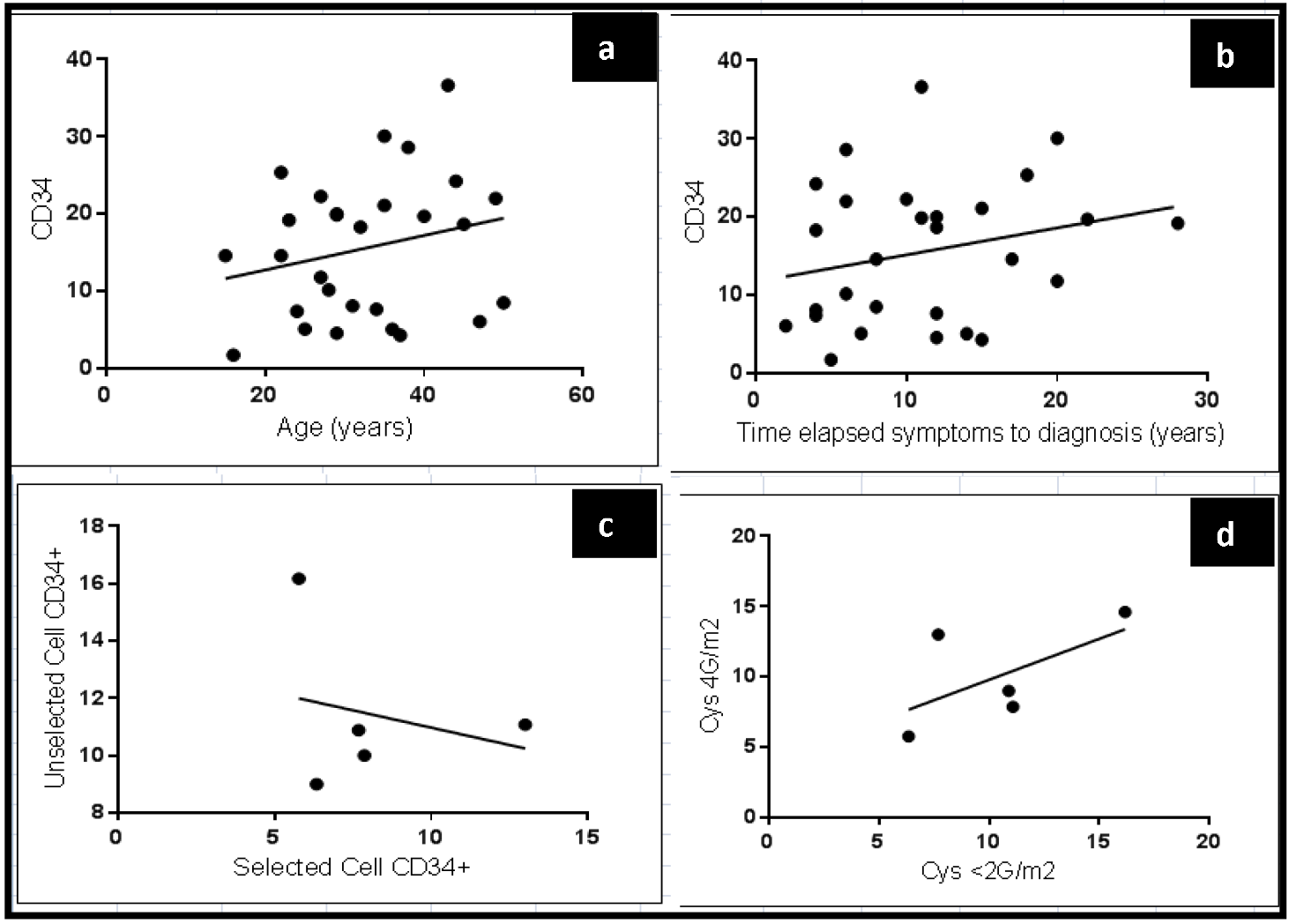
Pearson correlation analysis, a - comparison between harvested CD34+ cells and patients’ age at haematopoietic stem cell transplantation, b-comparison between elapsed time since onset of symptoms with the collection of PB-HPC. c - analysis between studies that manipulated the haematopoietic stem cell transplantation (selected) and those that did not (unselected) (Pearson r: 0.2492; two tailed p-value = 0.686). d - studies that used 2 or 4 g/m2 of cyclophosphamide during the mobilization regime for haematopoietic stem cell transplantation collection (Pearson r: 0.6038; two tailed p-value = 0.6038]

Table 3 lists the clinical events and the complications observed during the mobilization period.

**Table 3.**
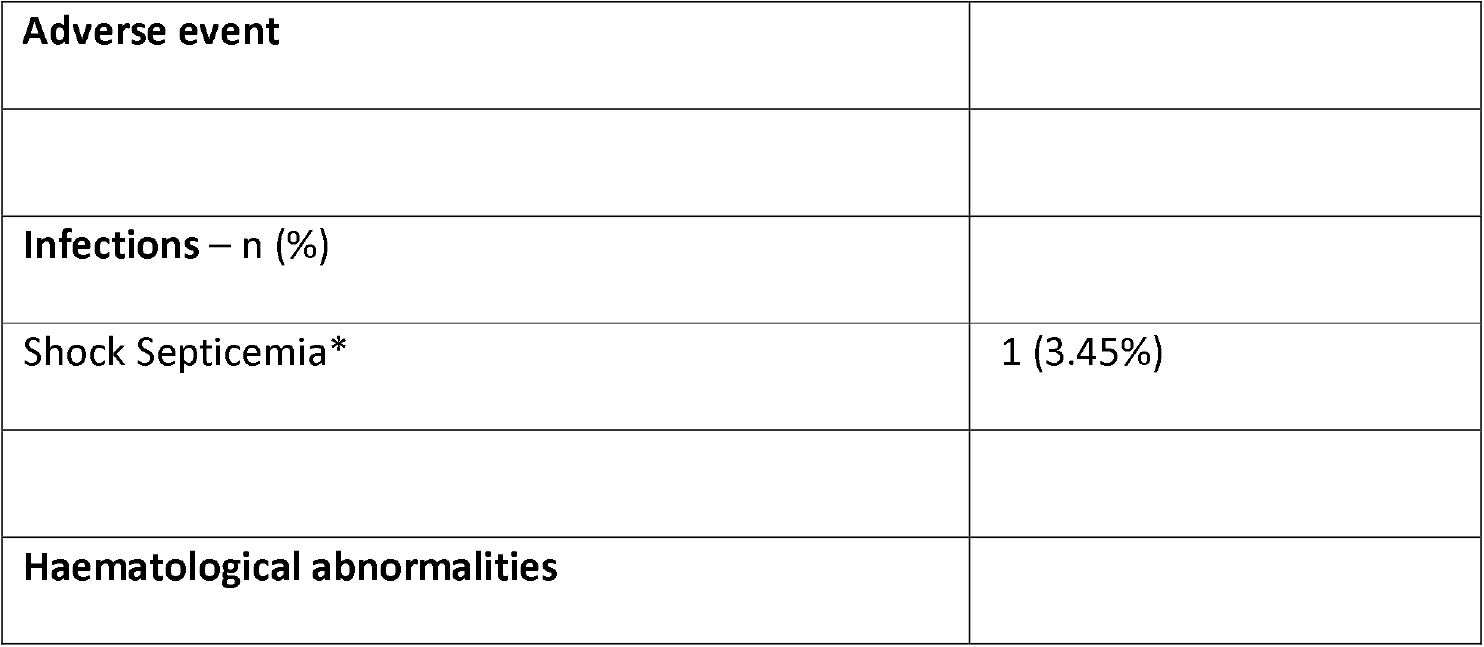
Complications and adverse events during the mobilization period.

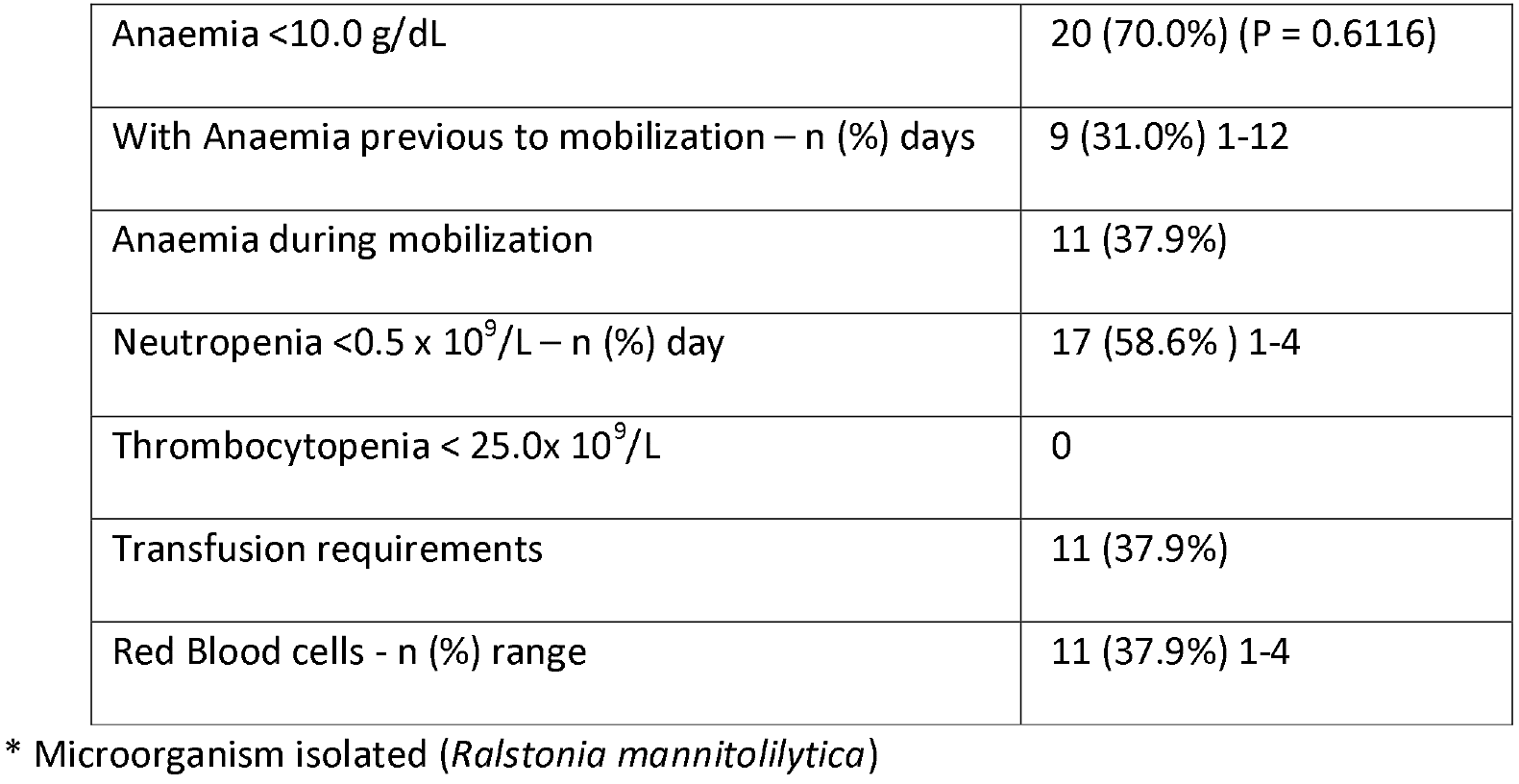

Table 4 lists the mobilization data from other autologous HSCT studies.

**Table 4.**
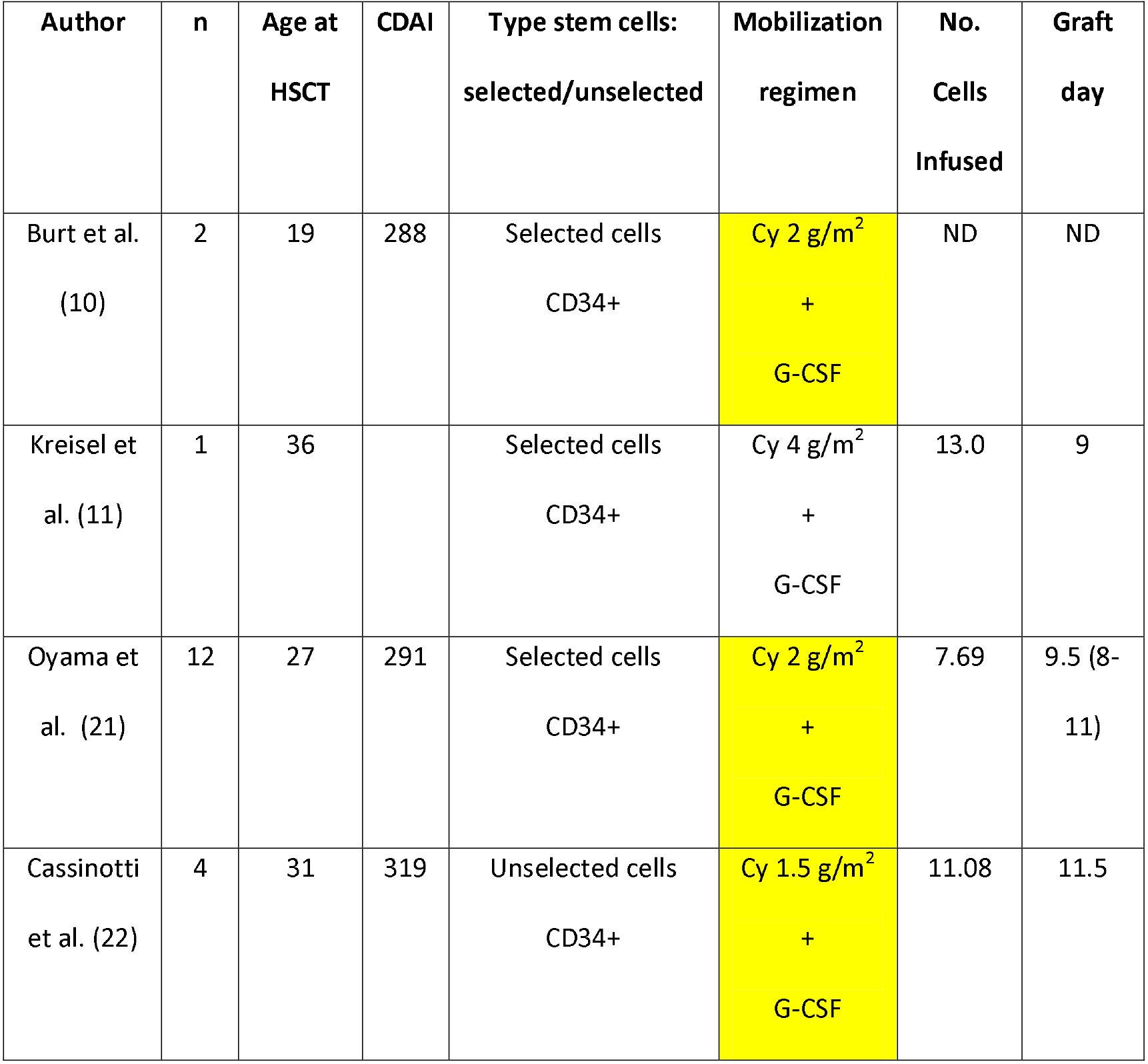
Previous studies of autologous haematopoietic stem cell transplantation (HSCT) for Crohn’s disease.

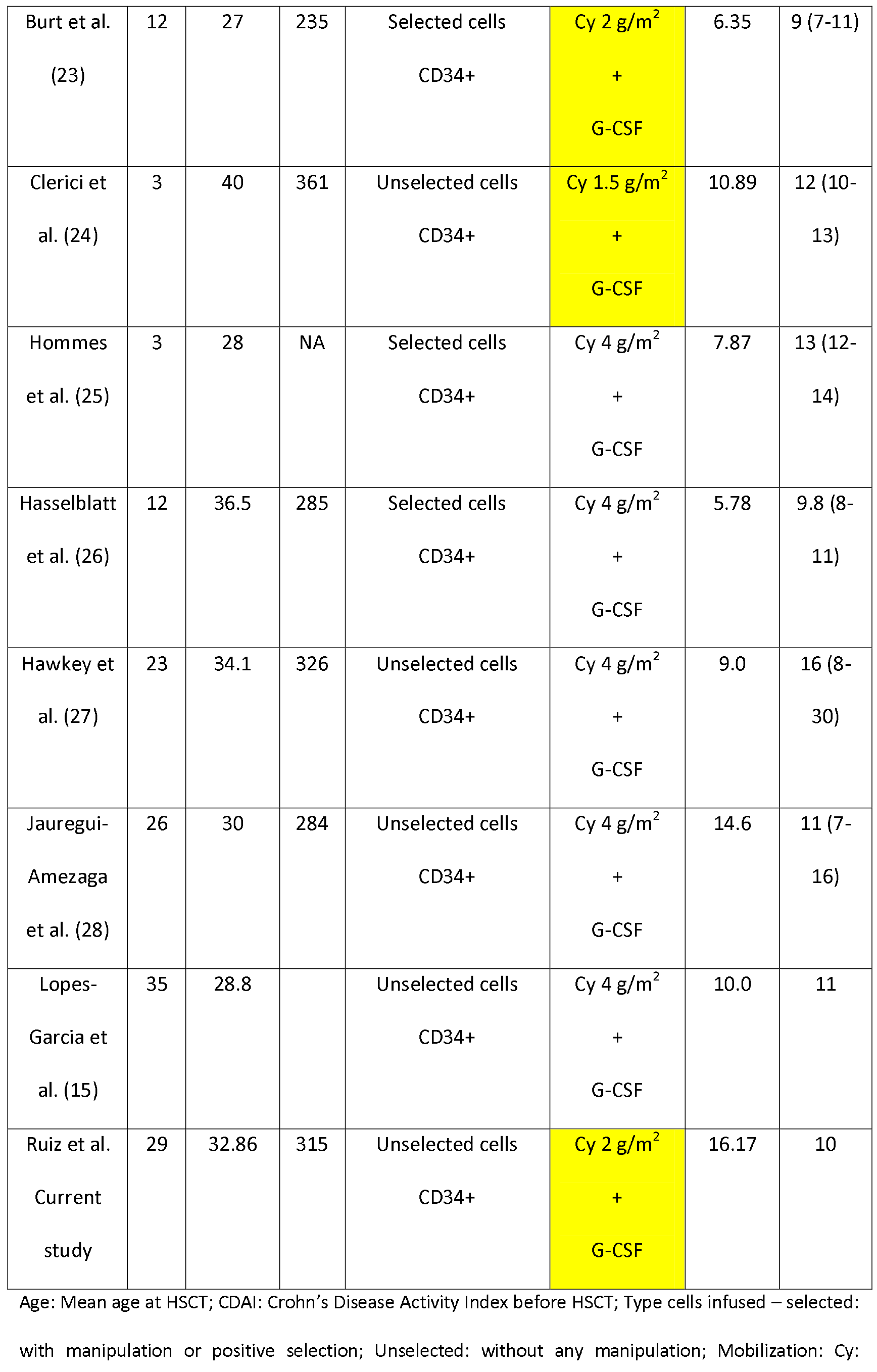

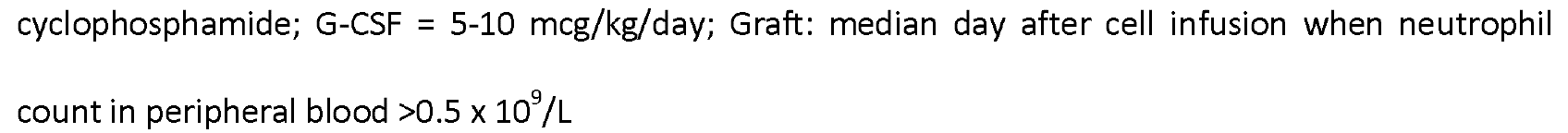

## Discussion

Autologous HSCT is a well-established procedure for the treatment of different hematologic malignancies and selected solid tumours and has become an option in the treatment of autoimmune diseases (29) (19). However, a number of concerns persist with regard to the mobilization regime (30). These issues include the choice of chemotherapy and administered doses, use of cytokines such as G-CSF and Granulocyte-macrophage colony-stimulating factor (GM-CSF) as well as the manipulation or not of the cells to be infused, the acceptable number of leukapheresis sessions and the criteria to begin harvesting the PB-HPC. Furthermore, new issues are arising such as the emergence of new agents such as the CXCL4 antagonist, AMD plerixafor, which is directed towards poor mobilizing patients and its pre-emptive use in cases of risk.

It is consensus that failure to mobilize is defined when a minimum number of 2.0 × 10^6^ CD34^+^ cells/Kg is not harvested in four leukapheresis sessions. The ultimate goal of collection for a safe transplant is 3.5 × 10^6^ CD34^+^ cells/Kg. When the minimum number of cells in the first session is not obtained, the patient must undergo further leukapheresis sessions in order to reach the target number. A marginal collection is considered when the number of collected cells in the first session is less than 3.0 × 10^6^ CD34^+^ cells/Kg.

Grafting is defined as the first day that the neutrophil count recovers to values greater than 0.5 × 10^9^/L in the PB with this number being maintained for two consecutive days.

These criteria have been supported by publications since the last century, including guidelines and the extensive literature that permeates the mobilization procedures for autologous and allogeneic HSCT in the treatment of malignant haematological diseases (31). This does not necessarily apply to HSCT for autoimmune diseases. The reports found are old and do not mention CD, the subject of this study.

The study by Burt et al. in 2001 was the first to address this subject by investigating data on stem cell collection of 187 patients from 24 centres in Asia, Australia, Europe and North America to determine the evolution, methods, effects on disease activity, and complications that might occur due to the procedure (32). Cases of multiple sclerosis predominated in the study, followed by rheumatoid arthritis, scleroderma and systemic lupus erythematosus. Few cases of other diseases have also been reported, but there is no mention of CD. It should be remembered that the first reports of HSCT specifically for CD appeared after 2003 (10) (11).

The investigation by Blanck et al., the second study to address this theme, evaluated 35 patients, 15 of whom had systemic sclerosis, 11 had multiple sclerosis and nine had other autoimmune diseases, and provided information on mobilization in these diseases (33) (30). Again, this study made no reference to CD (33).

Mobilization regimens use various chemotherapeutic agents including Cy and etoposide, or combinations of drugs such as dexamethasone, high-dose cytarabine and cisplatin (DHAP), or etoposide, methylprednisolone, cytarabine & cisplatin (ESHAP) with the aim of reducing tumour mass and improving the results of mobilization when the goal is to treat malignant haematological diseases.

Cy is the main chemotherapeutic agent used in mobilization regimes. Doses range from 2-7 g/m^2^ depending on the disease with great variability in toxicity. High doses of Cy cause high toxicity and controversial mobilization results (14).

In the first study, despite demonstrating a great variability in the regimens employed, the use of CY associated to G-CSF was shown to be the preferred regimen for mobilization in autoimmune diseases (32).

The second study confirmed this preference and reported that low doses of CY associated with multiple doses of G-CSF was effective in mobilizing PB-HPC (33). The dose of Cy was 4 g/m^2^ in 16 patients and 2 g/m^2^ in the others with similar efficiencies reported for both groups.

When the mobilization regimes for CD are observed, the rule is the use of Cy and G-CSF (Table 4).

The present study used the dose of 2 g/m^2^ of Cy. As in the other autoimmune diseases, we observed that there was no advantage in using the 4 g/m^2^ dose (33). It was confirmed that the dose of 4 g/m^2^ in addition to conferring greater toxicity to patients with CD (27) (34) does not provide benefits in the final counts of cells obtained (Table 4).

However, the question of the best mobilization regime for autoimmune diseases, including CD still lacks consensus (30). To support this statement, we highlight the recent proposal to reduce the dose of Cy to 1 g/m^2^ (35) as a result of warnings about toxicity in HSCT for CD (20) (34). The results of this study suggest that the reduction of Cy in the conditioning regime can be feasible without harming the result (20) (33) (30).

Another aspect regarding mobilization is related to the use of G-CSF. The literature describes differences in results depending on the origin of G-CSF (lenograstim, pegfilgrastim or filgrastim) (36). There are also questions about the varying doses used in different mobilization regimens that generally range from 5-10 mcg/kg/day but can occasionally go up to 16 mcg/kg/day. This higher dose apparently makes little difference to the final volume of cells collected (36). Hence, in all CD mobilization studies, the usual dose is 10 mcg/kg/day (in our case filgrastim) and always associated with Cy. This is due to reports of flares or exacerbation in patients with other types of autoimmune diseases submitted to HSCT (37). In respect to CD, there are no references or attempts to use G-CSF alone in mobilization, or the use of GM-CSF or an association with thrombopoietin as described in mobilization regimens for malignant haematological diseases (38).

The number of cells collected and made available for reinfusion is critical to the success of HSCT, as it reduces the period of aplasia and transfusion requirements, and promotes grafting. This was evident in our study, as well as in those described in Table 4, with no failure of mobilization or grafting occurring in any of the cases.

CD patients do not present harvesting difficulties similar to other autoimmune diseases such as multiple sclerosis (MS), rheumatoid arthritis (RA) and systemic lupus erythematosus (SLE), and in particular patients with systemic sclerosis where marginal collections or failures were observed due to intramedullary fibrosis and reduction of bone marrow microvasculature.(39)

On comparing the present work with other published studies, it appears that the composition of the cells infused during HSCT, whether manipulated or not, is unimportant to the outcome in respect to the day of grafting.

The rate of adverse events was low in our series, just one case of sepsis. Moreover, the transfusion requirements, restricted to packed red blood cell transfusions, was low despite the fact that most of the patients had anaemia throughout the mobilization period.

## Conclusion

Thus, we can conclude that CD patients are good mobilizers for HSCT. The dose of Cy should not exceed 2 g/m^2^ and it is necessary to have a dose-reduction study to see if the goal of reducing the dose further does not negatively affect the outcome.

